# Development of Scaffold-free 3D Cholangiocyte Organoids to Study the Progression of Primary Sclerosing Cholangitis

**DOI:** 10.1101/2022.10.24.513594

**Authors:** Wenjun Zhang, Konstantina Kyritsi, Abdulkadir Isidan, Yujin Park, Ping Li, Arthur A Cross-Najafi, Kevin Lopez, Lindsey Kennedy, Keisaku Sato, Shannon Glaser, Heather Francis, Gianfranco Alpini, Burcin Ekser

**Affiliations:** Division of Transplant Surgery, Department of Surgery, Indiana University School of Medicine, Indianapolis, IN; Division of Gastroenterology and Hepatology, Department of Medicine, Indiana University School of Medicine, and Division of Research, Richard L. Roudebush VA Medical Center, Indianapolis, IN; USA; Department of Medical Physiology, Texas A&M University College of Medicine, Bryan, Texas, USA

**Keywords:** 3D cell culture, cholangiocytes, human liver organoid, organoids, primary sclerosing cholangitis

## Abstract

Organoids are novel *in vitro* models to study intercellular crosstalk between the different types of cells in the pathophysiology of disease. To better understand the underlying mechanisms driving the progression of primary sclerosing cholangitis (PSC), we developed scaffold-free multi-cellular 3D cholangiocyte organoids (3D-CHO) using ‘primary’ liver cell lines derived from normal and PSC patients. Human liver samples from healthy donors and late-stage PSC patients were used to isolate ‘primary’ cholangiocytes (EPCAM^+^/CK-19^+^), liver endothelial cells (LECs, CD31^+^), and hepatic stellate cells (HSCs, CD31^−^/CD68^−^/Desmin^+^/Vitamin A^+^). 3D-CHOs were formed using cholangiocytes:HSCs:LECs and kept viable for up to 1 month. Isolated primary cell lines and 3D-CHOs were further characterized by immunofluorescence (IF), qRT-PCR, and transmission electron microscopy. Gene expressions for cholangiocytes (*SOX9, CFTR, EpCAM, AE, SCT, SCTR*), fibrosis (*ACTA2, COL1A1, DESMIN, TGFβ1*), angiogenesis (*PECAM, VEGF, CDH5, vWF*), and inflammation (*IL-6, TNF-α*) confirmed PSC phenotypes of 3D-CHOs. Since cholangiocytes develop a neuroendocrine phenotype and express neuromodulators, confocal-IF demonstrated that neurokinin-1 receptor (NK-1R, expressed by cholangiocytes and upregulated in PSC), was localized within CK-19^+^ cholangiocytes. Moreover, 3D-CHOs from PSC patients confirmed PSC phenotypes with upregulated NK-1R, tachykinin precursor 1, and downregulated membrane metalloendopeptidase. Our viable scaffold-free multiple-cell 3D-CHOs showed superiority as an *in vitro* model in mimicking PSC *in vivo* phenotypes compared to 2D cell culture, which can be used in PSC disease-related research.

## INTRODUCTION

Primary sclerosing cholangitis (PSC) is a heterogeneous and progressive liver disease mainly involving bile ducts with inflammation, ductular reaction, biliary senescence, and ultimately liver fibrosis.^1^ Patients with advanced stage of PSC often develop liver cirrhosis and may progress to the cholangiocarcinoma.^1^ There is currently no targeted therapy for PSC aside from supportive and immunosuppressive therapies.

Although small animal models, such as multidrug resistant-2-knockout (Mdr2)^-/-^ and bile-duct ligation (BDL) mouse models, help to study specific molecular pathways and identify new genetic drivers of PSC *in vivo*, they are limited by (i) lack of heterogeneity which occurs in patients with PSC and (ii) dissimilarities in transcriptomics and proteomics between humans and mice.^2^

Substance P (SP) is a neuropeptide that functions as a neurotransmitter and promotes ductular reaction and biliary senescence.^3^ We have demonstrated that PSC patients have significantly elevated serum levels of SP compared to healthy controls.^4^ SP binds to the neurokinin-1 receptor (NK-1R), and the expression levels of NK-1R are upregulated in liver tissues of PSC patients.^4^ Furthermore, inhibition of NK-1R by L-733,060 (NK-1R antagonist) attenuated ductular reaction, biliary senescence, and liver fibrosis in *Mdr2*^-/-^ (PSC model) mice, which suggests that the SP/NK-1R axis is a promising therapeutic target for the management of PSC.^4^

Organoids are a breakthrough in cell biology being tiny, self-organized, three-dimensional (3D) i*n vitro* cell cultures, which form a tissue construct that mimics its corresponding *in vivo* organ.^5^ Organoids can be generated by induced pluripotent stem cells, tissue pieces (e.g., liver biopsy), fluid/secretion (e.g., bile), or primary cells (e.g., cholangiocytes). When stem cells are used to generate cells, they are often referred to as “-like cells” (e.g., hepatocyte-like or cholangiocyte-like cells). There are several limitations of cell-like-derived organoids to model a liver disease; (i) they are often single lineage-cell derived (e.g., cholangiocyte-like-derived), (ii) they often use gel matrix to support the culture (e.g., Matrigel) which complicates the biology, transcriptomics, and proteomics since the most commonly used gel matrix is derived from mouse Engelbreth-Holm-Swarm sarcoma cells,^6,7^ and most importantly (iii) they often lack the 3D liver microenvironment due to the absence of support cells, such as hepatic stellate cells (HSCs) and liver endothelial cells (LECs).^8^ Previous *in vitro* human models for PSC included a 2D model of cholangiocytes isolated directly from PSC patients^9^ and 3D ‘cholangioids’ formed using human bile^10^, and cholangiocytes isolated from PSC livers.^11^ However, all previously reported 3D liver organoids or cholangioids lacked the human 3D *in vivo* disease microenvironment due to single lineage cell-derived 3D model.^8,11,12^

Here we demonstrate, for the first time, the development of scaffold-free multi-cell 3D cholangiocyte organoids (3D-CHO) formed by ‘primary’ human cholangiocytes, HSCs, and LECs isolated directly from PSC patients.

## MATERIALS AND METHODS

### Patient Samples

All PSC liver tissues were collected from explanted livers at the time of liver transplantation under a protocol approved by the Institutional Review Board of Indiana University School of Medicine (PI: Dr. Burcin Ekser). Multiple liver pieces (approximately 3cm^3^ each) were harvested from the right, left, and caudate lobes of the same liver for cell isolation to include heterogeneity of the disease (Figures 1A and 1B). Moreover, extrahepatic bile ducts were also collected for cholangiocyte isolation to include another heterogeneity of cholangiocytes, such as intrahepatic and extrahepatic cholangiocytes. Although explanted PSC livers represented late-stage PSC, all livers were not similarly cirrhotic in our clinical observation due to the disease progression and extra model for end-stage liver disease points granted to certain patients for clinical symptoms before transplantation so that they could be moved higher in the waitlist to prioritize liver transplantation (Figures 1A, 1B, and 1C). All normal liver tissues were collected during organ procurement surgeries approved by the Indiana Donor Network (PI: Dr. Burcin Ekser) (Figure 1C, right panel). Patient characteristics for PSC and normal liver tissues are provided in Table 1.

**Figure 1:**
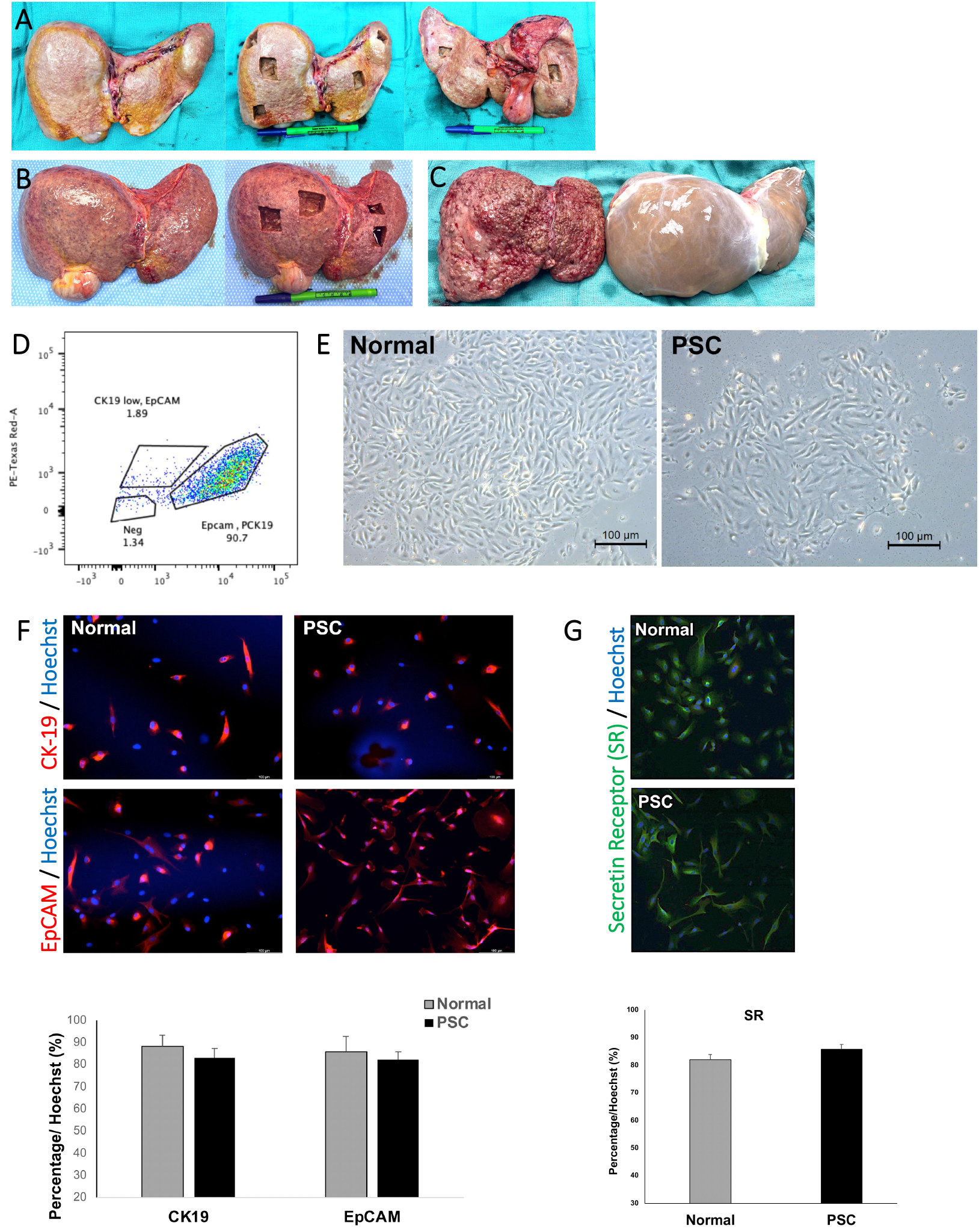
Isolation and characterization of cholangiocytes from normal and PSC patients. **A**. A representative picture of an explanted PSC liver and how liver samples (∼3 cm^3^ / each) were obtained from right, left, and caudate lobes for cell isolation. All pictures (right, middle, and left panels) show the same liver before and after the liver samples were obtained. **B**. Another representative picture of an explanted PSC liver at the time of liver transplantation shows a different level of cirrhosis. Both the right and left panels show the same liver. **C**. Representative pictures of an explanted PSC liver with dense fibrosis and advanced cirrhosis (left panel) as well as the normal liver (right panel). **D**. A representative FACS analysis for cholangiocyte isolation from a female normal donor. The EpCAM^+^CK-19^+^ cholangiocytes were over 90% of all cell populations. **E**. Brightfield image of cultured human normal and PSC cholangiocytes after FACS purification. **F**. IF staining of CK-19 (top, in red) and EpCAM (below, in red) of human immortalized normal (left panel) and PSC (right panel) cholangiocytes. Hoechst (blue) staining for nuclear counterstain. Lower, quantification of the percentage of CK-19^+^ and EpCAM^+^ cell (% Hoechst stained nucleus) in isolated normal and PSC cholangiocytes show no significant difference in CK-19 and EpCAM staining between normal and PSC cholangiocytes (n=3 patients for normal and PSC) **G**. IF staining of human immortalized Normal (top) and PSC (bottom) cholangiocytes. SR (secretin receptor) staining (green) for cholangiocytes and Hoechst (blue) staining for nucleus counterstain. Bottom, quantification of SR^+^ cells percentage (% Hoechst stained nucleus) in isolated normal and PSC cholangiocytes show no significant difference in SR staining between normal and PSC cholangiocytes. n=3 patients for normal and PSC. Scale bars: 100µm.

**Table 1:** Patient characteristics for liver cell isolation from normal and primary sclerosing cholangitis.

| Patient # | Age (years) | Gender | Blood Type | Diagnosis |
| --- | --- | --- | --- | --- |
| 40 | 24 | M | A | Donor |
| 48 | 18 | M | A | Donor |
| 65 | 46 | M | O | Donor |
| 68 | 23 | F | A | Donor |
| 71 | 39 | F | B | Donor |
| 77 | 29 | M | A | Donor |
| 83 | 38 | M | A | Donor |
| 93 | 19 | M | B | Donor |
| 97 | 42 | M | AB | Donor |
| 105 | 39 | M | O | Donor |
| 115 | 35 | M | A | Donor |
| 116 | 43 | M | O | Donor |
| 61 | 70 | F | O | PSC |
| 62 | 61 | M | O | PSC |
| 67 | 38 | M | O | PSC |
| 76 | 56 | F | O | PSC |
| 85 | 46 | F | A | PSC |
| 89 | 55 | M | O | PSC |
| 92 | 60 | M | A | PSC |
| 99 | 32 | M | O | PSC |
| 118 | 40 | F | B | PSC |

**Legend:** M= male, F= female, PSC = primary sclerosing cholangitis. Cells isolated from normal and PSC patients were not used in the order indicated above. Different cells were used for different assays. Organoids formed by normal and PSC cells were not compared to each other one by one, but they were grouped as indicated in the manuscript.

### Isolation and Maintenance of Liver Cells

For isolation of primary human cholangiocytes, LECs, and HSCs, the explanted liver tissues were washed with 1x phosphate-buffered saline (PBS) solution to remove residual red blood cells. The washed liver tissues were minced into small pieces (<3 mm^3^) and then incubated with the cell dissociation buffer composed of Eagle’s Minimum Essential Medium (EMEM) supplemented with a mixture of collagenase type XI and DNase (Sigma-Aldrich, St. Louis, MO) at a concentration of 250 mg/L and 50 mg/L, respectively, for 20-30 minutes, depending on the severity of stiffness due to liver cirrhosis in each liver.

Following tissue digestion, liver non-parenchymal cells (NPCs) and cholangiocytes were enriched by three consecutive centrifugation steps at 100x *g* for 3 minutes each at 4°C. After three washes with EMEM, the cell pellet was suspended in a seeding medium and cultured on collagen-coated plates. The isolated intrahepatic and extrahepatic cholangiocytes were enriched with flow cytometry (FACS) sorting using epithelial cellular adhesion molecule (EpCAM) antibody (Progen, Heidelberg, Germany, Catalog No. 61104) for EpCAM^+^ cells. The enriched cells were further confirmed by immunostaining for cytokeratin-19 (CK-19) positivity (Abcam, Boston, MA, Catalog No. ab7754) (Figure 1D), and subsequently cultured in a mixture of Dulbecco’s Modified Eagle Medium (DMEM) (Gibco, Waltham, MA) and DMEM/F12 medium (1:1), supplemented with 10% fetal bovine serum (FBS), penicillin-streptomycin (1x), adenine (100µM), insulin (5µg/ml), epinephrine (1µg/ml), T3-T(8µg/ml for transferrin), epidermal growth factor (EGF) (10ng/ml), and hydrocortisone (600ng/ml). The NPCs that remained in the supernatant were centrifuged at 600x*g* for 10 minutes at 4°C, and then resuspended to purify LECs with FACS to select CD31^+^ cells (Miltenyi Biotech, Bergisch Gladbach, Germany, Catalog No. Cat# 130-117-312). LECs were cultured in EGM™-2 endothelial growth medium (Lonza, Basel, Switzerland). Primary HSCs were collected after selecting Kupffer cells and LECs, as previously described.^13^ HSCs were maintained in low glucose DMEM (Gibco, Waltham, MA) medium supplemented by FBS (10%), penicillin-streptomycin (1x), and Glutamax (1x, Gibco). For long-term cell culturing and organoid-creating purposes, the isolated normal and PSC cholangiocytes, HSCs, and LECs were also immortalized with HPV-16 E6/E7 expressing lentivirus (ABM, Richmond, BC).

### 3D Cholangiocyte Organoid Formation

We tested different combinations of cholangiocytes for 3D-CHO formation, such as (i) cholangiocytes alone, (ii) cholangiocytes and LECs with a 2:1 ratio, (iii) cholangiocytes and HSCs with 2:1 ratio, and (iv) cholangiocytes, HSCs, and LECs with 2:1:1 ratio. A total of 40,000 cells from primary as well as immortalized normal and PSC liver cells (cholangiocytes, LECs, and HSCs) were mixed at the above-indicated ratios in the organoid culturing medium (H69 medium: HSC medium: LEC medium in 2:1:1 ratio), and then the cell mixture was cultured in 96 well U-bottom plates with an ultralow attachment surface (S-Bio, Hudson, NH) for two days to form the cell aggregate. 3D-CHOs were able to be maintained in the medium for up to 30 days without significant loss of cell viability. The morphology was similar in all established organoids regardless of cell types included (Figure 3B), indicating the feasibility of 3D-CHOs formation with the scaffold-free method.

### RNA Extraction and qRT-PCR

Total RNA extraction and isolation from normal and PSC liver cells or 3D-CHOs were performed using Trizol™ reagent and a PureLink RNA kit (Invitrogen, Waltham, MA). cDNAs were synthesized with a SuperScript™ First Strand Synthesis System (Invitrogen). Quantitative PCR (qRT-PCR) was performed using SsoAdvanced Universal SYBR Green Supermix (Bio-Rad, Hercules, CA, Catalog No. 1725271). The relative expression level was normalized to the housekeeping gene HPRT1 or GAPDH. The mRNA expression levels of each gene were calculated with the 2^−ΔΔCT^ method. Cholangiocyte marker genes (*SOX9, EPCAM, CFTR, AE2, SCT, SCTR*), fibrosis marker genes related to HSCs (*ACTA2, COL1A1, Desmin, TGFβ1*), angiogenesis marker genes (*PECAM, CDH5, vWF, VEGF*), and inflammation marker genes (*IL-6, TNFα)* were measured and compared between normal and PSC 3D-CHOs. Moreover, the gene expression of NK-1R, tachykinin precursor 1 (TAC1), and membrane metalloendopeptidase (MME) was also measured and compared between normal and PSC 3D-CHOs. The primers used are listed in Table 2.

**Table 2:**
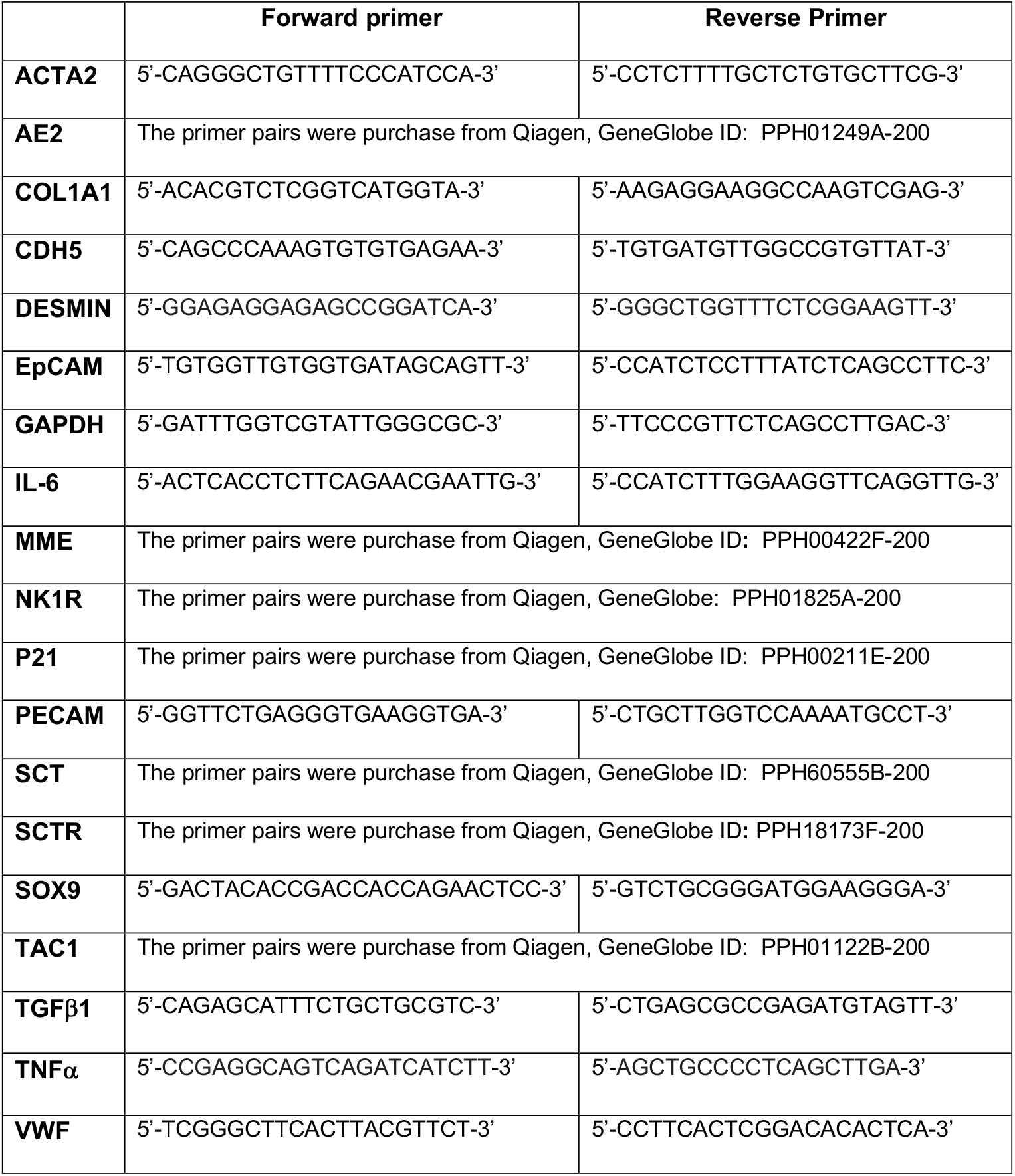
List of human PCR Primers.

### Flow cytometry, Immunofluorescence (IF) Staining, and Transmission Electron Microscopy (TEM)

For FACS of EpCAM and CK-19 positive cell population in purified and immortalized normal and PSC cholangiocytes, the cells were fixed and permeabilized with FluoroFix solution (Biolegend) and intracellular staining perm wash buffer (Biolegend), as instructed by the manufacturer. The fixed and permeabilized cells were then incubated with primary antibodies against CK-19 (Rabbit, Abcam, Boston, MA, Catalog No. ab52625) and EpCAM (mAb Progen, Heidelberg, Germany, Catalog No. 61104), and followed by incubation with fluorescent dye conjugated secondary antibodies (ThermoFisher Sci) before being subjected to FACS analysis using BD LSRFortessa flow cytometry analyzer.

For IF co-staining, the organoids were fixed in 2% paraformaldehyde for 15 minutes and permeabilized with 0.1% Triton X-100 for 30 minutes. Blocking was performed with PBS containing 0.05% Tween and 5% normal goat serum for 60 minutes. Organoids were then incubated with primary antibodies overnight at 4°C, followed by incubation with the appropriate secondary antibodies for 1 hour at room temperature. Images were captured with a Leica DMi8 Microscope and Imaging System (Wetzlar, Germany). Antibodies used for IF staining are: CK-19 (Abcam, catalog No. ab7754), CK-7 (ThermoFisher Sci, catalog No. MA1-06315), Secretin receptor (SR) (Abcam, catalog No. ab234830), and cystic fibrosis transmembrane conductance regulator (CFTR) (Abcam, catalog No. ab2784) for cholangiocytes, vascular endothelial-cadherin (VE-Cadherin) (ThermoFisher Sci, catalog No.14-1449-82) and CD31 (Milteny Biotech, catalog No.130-117-390) for LECs, and Desmin (Sigma, catalog No. 243R), Collagen Type 1 alpha 1 (COL1A1) (ThermoFisher Sci, catalog No. PA5-29569), smooth muscle α-actin (αSMA) (Sigma, catalog No. A2547), Periostin (POSTN) (R&R system, catalog No. MAB3548) and glial fibrillary acidic protein (GFAP) (ThermoFisher Sci, catalog No. PAI-9565) for HSC. Cell nuclei were countered with Hoechst 33342 (ThermoFisher Sci). The antibody against NK-1R was purchased from Novus (catalog No. NB100-74469). The images were taken by the Leica Immunofluorescence Microscope and digital camera system (Weltzar, Germany). The automated counting of single-color cell images was performed using ImageJ (U. S. National Institutes of Health, Bethesda, Maryland, USA, https://imagej.nih.gov/ij/) to quantify cells with fluorescence intensity reaching the threshold. Three random individual images were analyzed.

For TEM, the cholangiocyte organoids were fixed with TEM fixative containing 2% paraformaldehyde/2.5 glutaraldehyde in 100mM cacodylate buffer (Electron Microscopy Sciences, Hartfield, PA) for 2 hours at room temperature. The tissue processing, embedding, and TEM imaging were performed at the Department of Molecular Microbiology, Washington University School of Medicine.

### Measurement of Rhodamine-123 (Rh123) in 3D-CHOs

The measurement of Rh123 in organoids was performed as previously described^14,15^ with minor modifications. Briefly, the organoids on day 6 were divided into 2 groups, in which the organoids were preincubated with dimethyl sulfoxide (DMSO) (0.05%) or P-glycoprotein (P-gp) inhibitor, verapamil (20µM) (MedChemExpress, catalog No. HY-14275), for 15 minutes at 37°C before incubating with Rh123 (10µM) (Sigma, catalog No. R-8004) for another 15 minutes.

Then, 3D-CHOs were collected and washed twice in PBS before homogenization. The homogenate was then centrifuged at 12,000 × *g* for 5 mins at 4°C to collect the supernatant and read the fluorescence intensity (λ_ex_/λ_em_ = 485/520 nm) by Synergy H1 Hybrid Reader (Agilent, Vermont). The accumulation of Rh123 in 3D-CHOs was also visualized and imaged under fluorescence microscopy after incubation with Rh123 and verapamil.

### Neurokinin-1 Receptor (NK-1R)

Since cholangiocytes develop a neuroendocrine phenotype and express neuromodulators, the expression of NK-1R was checked by immunohistochemistry (IHC) in normal and PSC liver tissues, by IF in primary human cholangiocytes isolated from normal and PSC livers, as well as in 3D-CHOs in order to confirm the same phenotype of cells from the parent tissue, isolated in 2D culture and in 3D organoid formation. Moreover, the co-localization of NK-1R with cholangiocyte and HSC markers was also determined by IF in primary 2D cholangiocytes and HSCs, as well as in 3D-CHOs.

### Statistical Analysis

Differences between experimental groups were analyzed by the Student’s unpaired t-test (if two groups) or by one-way ANOVA (if more than two groups). Data are expressed as mean +/-SEM. A p-value of <0.05 was considered statistically significant.

## RESULTS

### Normal and PSC Hepatic Cell Isolation, Characterization, and Immortalization

The resected human liver tissues from normal donors and PSC patients were dissociated using Type IV collagenase and DNase I for human cholangiocytes, LECs, and HSCs isolation. Due to the different cirrhotic natures of each PSC liver, the application time of collagenase varied among PSC livers, as indicated above (Figures 1A, 1B, 1C). Purified and immortalized normal and PSC cholangiocytes showed a typical epithelial cell morphology (Figure 1E). FACS analysis (Figure 1D) and IF staining further confirmed that more than 80% (% Hoechst+ nuclei) of cholangiocytes were positive for the expression of EpCAM (normal = 85.09% vs PSC = 82.02%), CK-19 (normal = 88.09% vs PSC = 83.12%), and SR (normal = 82.05% vs PSC = 85.67%) [17] (Figure 1F and 1G).

Purified and immortalized HSCs and LECs from normal donor and PSC patients were further characterized by IF staining. We found that while the percentage of Desmin^+^ cells was comparable between HSCs from normal and PSC patients (Figure 2A), the ratio of positive cells for activated HSC markers, such as αSMA, POSTN, and GFAP, was significantly higher in HSCs from PSC patients compared to normal patients, confirming PSC phenotypes (Figures 2A and 2B). IF staining also showed that around 90% of LECs isolated from normal and PSC liver explants were positive for CD31 (Figure 2C). Consistently, the qRT-PCR analysis demonstrated that HSCs were specifically positive for Desmin and LECs were specifically positive for PECAM (CD31) in both normal and PSC patients (Figure 2D), confirming again their cellular characteristics.

**Figure 2:**
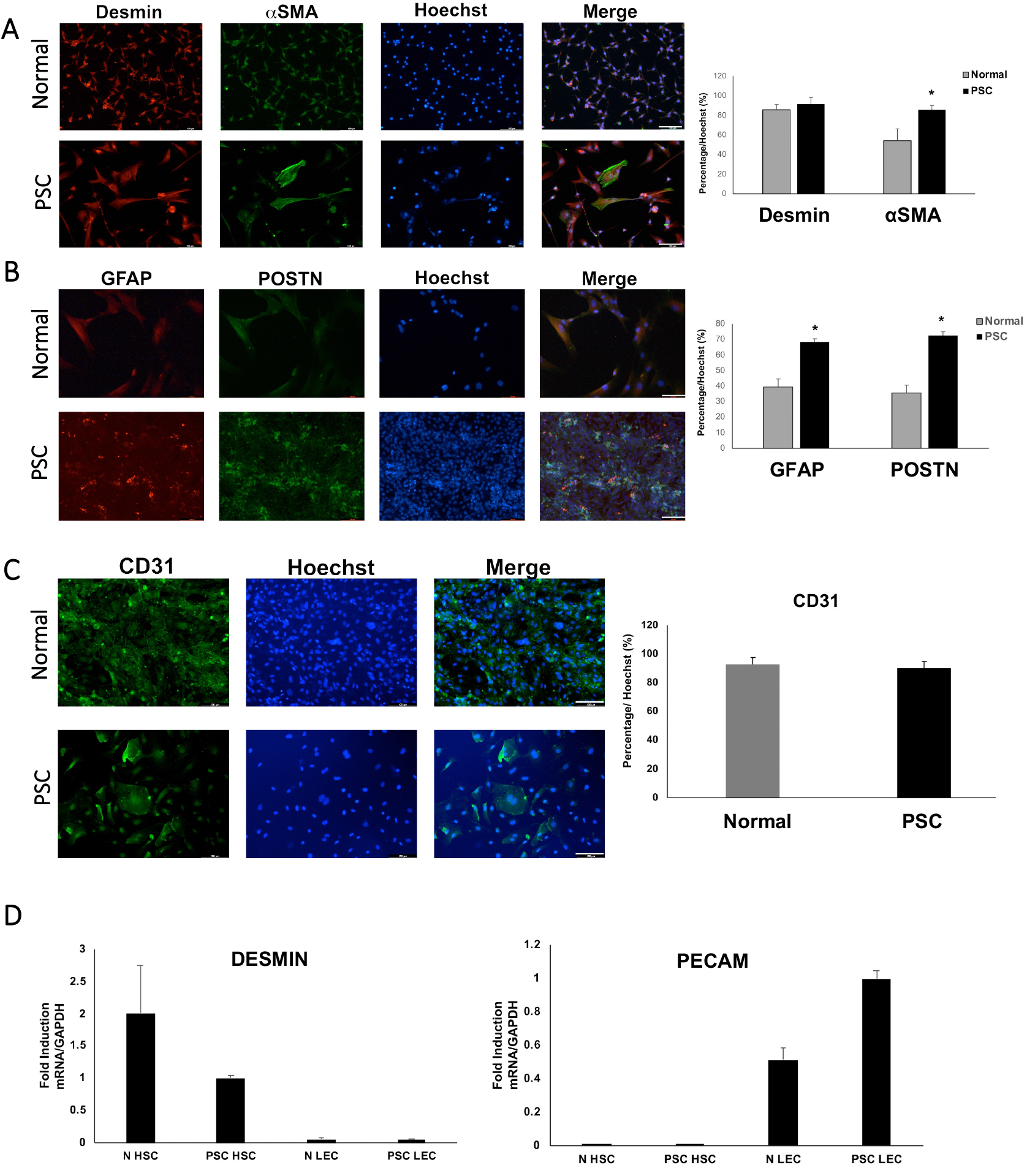
Isolation and characterization of HSCs and LECs from normal and PSC patients. **A**. IF staining for Desmin (red), and αSMA (green) of immortalized human HSCs from normal and PSC patients. Right, quantification of Desmin^+^ and αSMA^+^ cell percentage (% Hoechst stained nucleus) in isolated normal and PSC HSCs (n=3 patients for normal and PSC) (* = p <0.05). **B**. IF staining for GFAP (red), and POSTN (green) of immortalized human HSCs from normal and PSC patients. Right, quantification of GFAP^+^ and POSTN^+^ cells (% Hoechst stained nucleus) in isolated normal and PSC HSCs (n=3 patients for normal and PSC) (* = p <0.05). **C**. IF staining for CD31 (green) of immortalized human LECs from normal and PSC patients. Hoechst (blue) staining for nucleus counterstain. Right, quantification of CD31^+^ cells (% Hoechst stained nucleus) in isolated normal and PSC LECs. Scale bars: 100µm. **D**. qRT-PCR analysis of the expression of Desmin and PECAM in HSCs and LECs from normal and PSC patients, confirming cell phenotypes and characteristics.

### Generation of Scaffold-Free 3D Cholangiocyte Organoids (3D-CHO)

We tested different combinations of hepatic cells to create scaffold-free 3D-CHOs and all different cell combinations formed uniform 3D-CHOs with a diameter of around 150 micrometers on day 3 (Figure 3B). TEM demonstrated that tight junctions formed between cholangiocytes in the 3-cell 3D-CHOs as early as day 7 and peaked at day 14 (Figure 3C). Interestingly, 3D-CHOs composed of all 3 cells (cholangiocytes+LECs+HSCs) formed the typical ring-like ductal structure within the organoid and the surrounding cell aggregate area at 24 to 48 hours after organoid formation (Figure 3D). To verify the functionality of 3D-CHOs, the function of P-glycoprotein (P-gp) was determined by incubating 3D-CHOs with the fluorescent dye rhodamine 123 (Rh123). Rh123 was chosen as a fluorescent dye substrate because it could be sensitively visualized and detected by fluorescence microscopy and an automatic microplate reader.^16,17^ Both the fluorescence microscopy (Figure 3E, left panel) and an automatic microplate reader that examined the fluorescence (λ_ex_/λ_em_ = 485/520 nm) in the lysate of 3D-CHOs (Figure 3E,right panel) confirmed that blocking the transport activity with verapamil resulted in the accumulation of the Rh123 in the cytoplasm in the cells, which confirm the specific of P-gp mediated transport activity in 3D-CHOs. By day 4, the ring-like ductal structures were fully integrated into the cholangiocyte organoids, and IF staining demonstrated the CK-7 positive cells were distributed mainly in the center of the organoids, VE-cadherin positive LECs and αSMA positive HSCs presented in both the central and peripheral regions of the organoids (Figure 3F).

**Figure 3:**
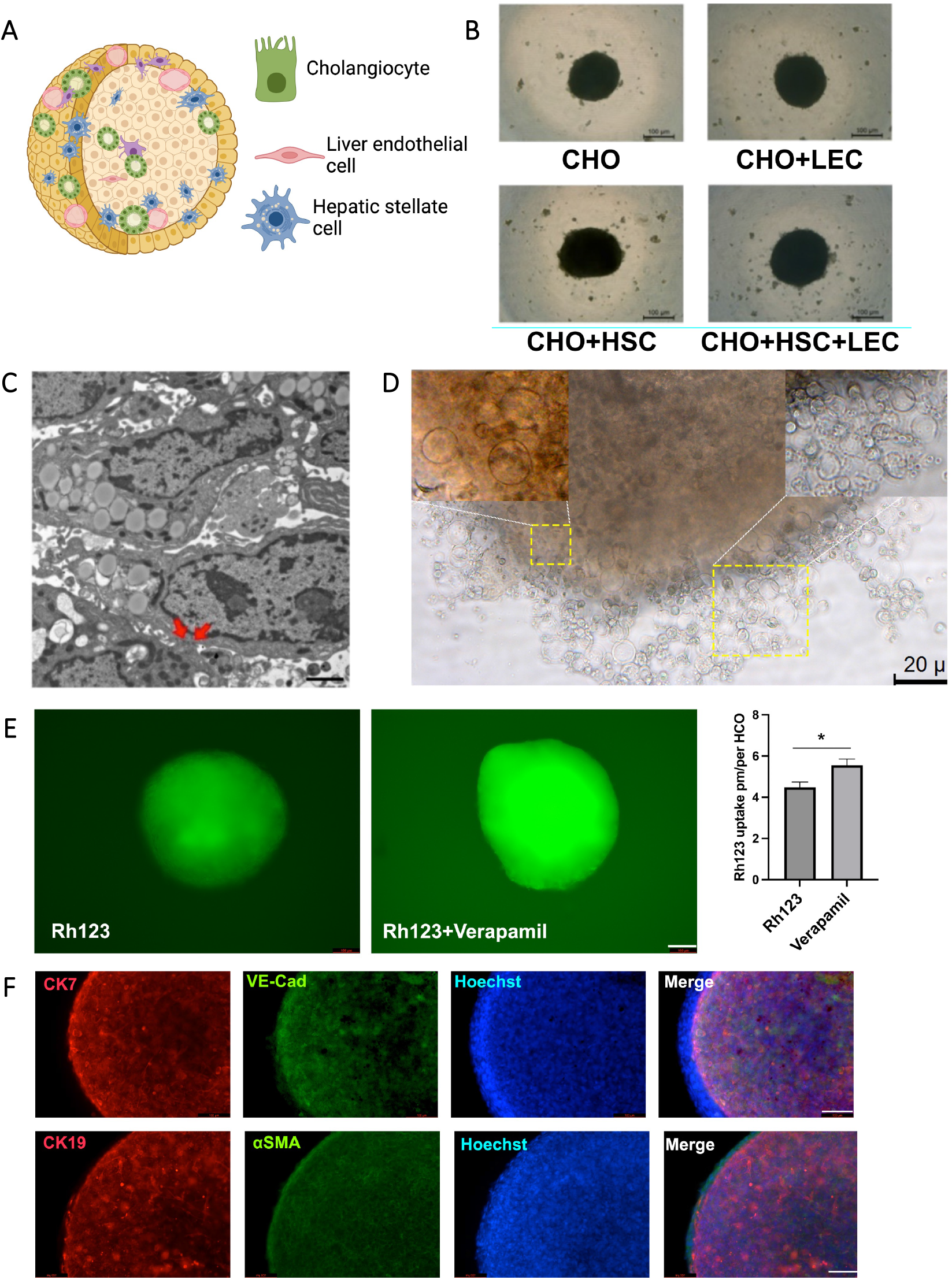
Scaffold-free cholangiocyte organoid formation. **A**. Representative cartoon of how multiple-cell 3D cholangiocyte organoids are formed with cholangiocytes (CHOs), hepatic stellate cells (HSCs), and liver endothelial cells (LECs) (created using biorender.com). **B**. Representative brightfield images of cholangiocyte organoids established using CHO, HSC, and LEC. A total of 20,000-40,000 cells are used to form organoids with a different ratio, such as CHO: HSC: LEC = 2:1:1. **C**. TEM image of cholangiocyte organoids in 3 cells. Cholangiocytes interact with each other and form 3D architecture *in vitro*. Red arrows indicate cell junctions (tight junctions) between cholangiocytes. Scale bar: 100 nm. **D**. The area with ring-like ductal structure within and outside of the organoid are amplified and indicated in the left and right corner of the image respectively. Scale bar: 20 µm. **E**. Left, fluorescence microscopy visualization of Rh123 in 3D-CHOs with or without preincubation of verapamil. Right, automatic microplate reader that examines the Rh123 fluorescence (λ_ex_/λ_em_ = 485/520 nm) in the lysate of 3D-CHOs (* = p<0.05) (n= 4 organoids for each experiment). **F**. Upper, representative IF staining image for CK7 (in red for cholangiocytes) and vascular endothelial cadherin (VE-Cad) (in green for LECs) in 3D-CHOs. Lower, representative IF image for CK19 (in red for cholangiocytes) and αSMA (in green for HSCs) in 3D-CHOs. The cell nucleus was stained with Hoechst (in blue) for cholangiocytes, LECs, and HSCs. Peripheral cells that are negative for VE-Cad and CK-7 are HSCs. The accurate amplification of the image is 100x.

### PSC Cholangiocytes Enhanced Angiogenesis and HSC Proliferation in Cholangiocyte Organoids

We next generated the 3-cell 3D-CHOs using immortalized cholangiocytes, LECs, and HSCs (cell ratio as 2:1:1) derived from normal donors and PSC patients in order to verify the ability of immortalized cells to mimic similar phenotypes as primary cells from PSC patients. Unlike normal 3D-CHOs that showed a uniform size and organoid shape by day 4, 3D-CHOs derived from the PSC patient showed a disorganized shape with the formation of detached cell aggregates surrounding the main organoids (Figure 4A). CellTiter-Glo 3D assay demonstrated that 3-cell PSC cholangiocyte organoids showed higher cell proliferation than normal 3-cell cholangiocyte organoids despite their formation with immortalized cells. This finding is consistent with ductular reaction (cholangiocyte proliferation) that is observed in PSC patients which confirms the PSC phenotypes in 3D-CHOs (Figure 4B).

**Figure 4:**
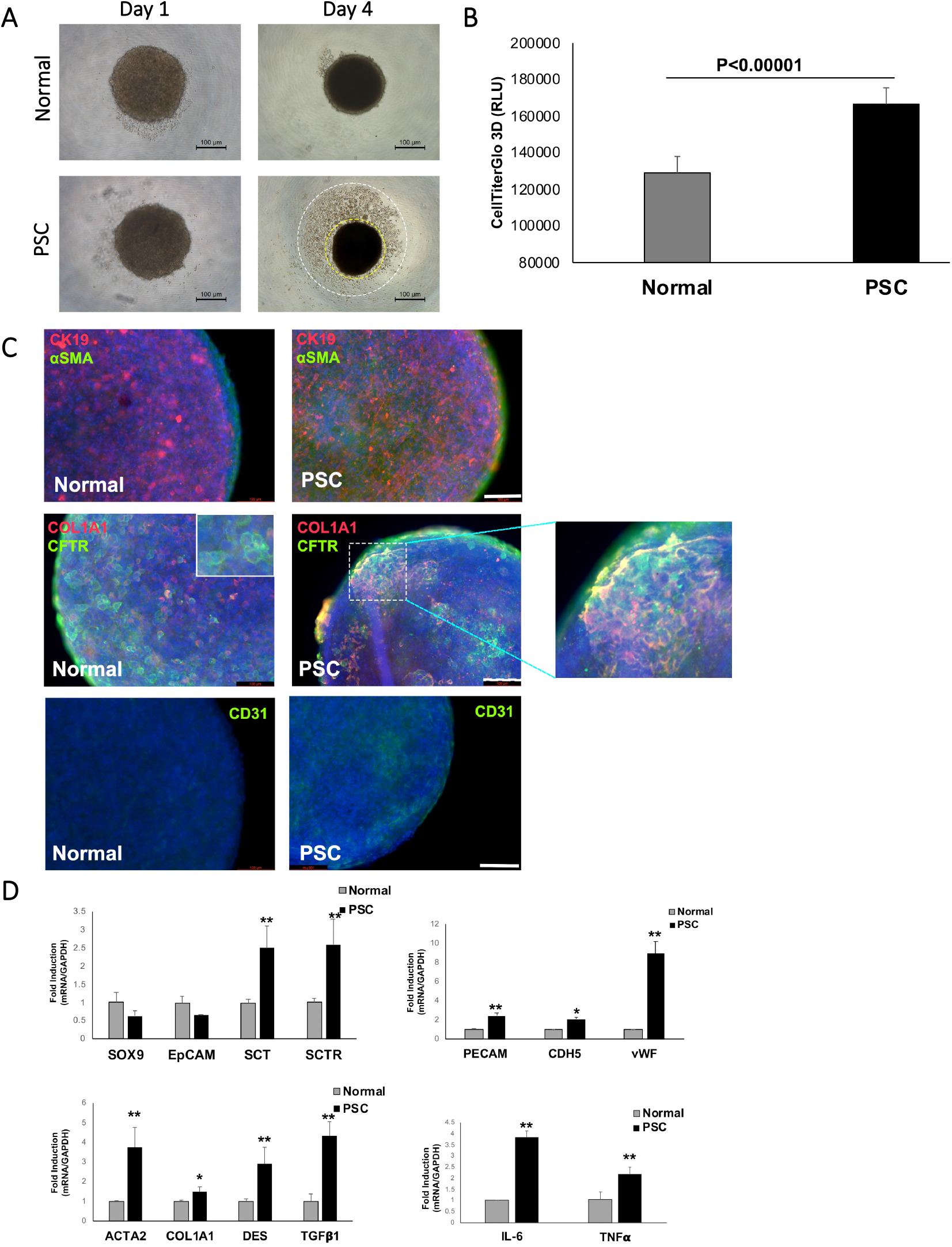
Gene expression in scaffold-free cholangiocyte organoids from normal and PSC patients. **A**. Representative brightfield images of 3-cell 3D-CHOs (cholangiocytes, LECs, HSCs) from normal and PSC patients on day 1 and day 4. A large number of cell aggregates (within the inner white line circle and outer layer circle) detached from the main organoid (within the inner circle) were observed in PSC organoids on day 4. **B**. CellTiterGlo 3D assay on normal and PSC 3D-CHOs showing significant proliferation in PSC cells after culturing for 5 days (n=3 per normal and PSC organoids). **C**. Top: representative IF co-staining image for CK-19 (in red for cholangiocytes) and αSMA (in green for HSCs) in normal and PSC organoids. Middle: representative IF co-staining image for CFTR (in green for cholangiocytes) and COL1A1 (in red for HSCs) in normal and PSC organoids. The side magnified image shows a nodular-like image in PSC organoids. Bottom: representative IF staining image for CD31 (in green for LEC vascularization) in normal and PSC organoids. Cell nucleus was stained with Hoechst (in blue). Scale bars: 100µm. **D**. qRT-PCRs for cholangiocyte marker genes Sox9, EpCAM, SCT (secretin) and SCTR (secretin receptor), fibrosis marker genes ACTA2, COL1A1, DESMIN (DES), and TGFβ1, angiogenesis marker genes PECAM, CDH5, and vWF, and inflammation marker genes IL-6 and TNFα. *P<0.05, **p<0.01 PSC vs. normal organoids. (n=4 organoids per each experiment).

IF staining further confirmed the PSC phenotypes in 3D-CHOs from PSC patients which had an increased expression of αSMA and COL1A1 compared to normal cholangiocyte organoids (Figure 4C). Interestingly, IF staining of CFTR and COL1A1 revealed that CFTR^+^ cholangiocytes were distributed evenly in normal 3D-CHOs without co-localization with COL1A1; however, in PSC 3D-CHOs, CFTR^+^ cholangiocytes were observed to form cell aggregates and surrounded with COL1A1 similar to nodular structures seen in cirrhotic livers (Figure 4C). In addition, PSC organoids had an elevated CD31 expression compared to normal 3D-CHOs (Figure 4C). The qRT-PCR analysis further demonstrated that, compared to normal organoids, PSC organoids have (i) an elevated expression of secretin and SR, (ii) increased expression of fibrosis maker genes, Acta2 (encodes αSMA), COL1A1, desmin, and TGFβ1, (iii) increased expression of angiogenesis marker genes PECAM, CDH5 (encodes VE-Cadherin), and von Willebrand Factor (vWF), and (iv) increased expression of inflammatory marker genes, such as IL-6 and TNFα, compared to normal organoids (consistent with IF staining results). Taken together, these results indicate that there is HSC activation and angiogenesis in PSC organoids (Figure 4D). These findings suggested that our 3-cell scaffold-free 3D-CHOs mimic the phenotypes and the pathophysiological microenvironment of the liver portal area found in PSC patients.

### 3D-CHOs from PSC Patients Maintain Neuroendocrine Phenotypes for up to 1 month

Our previous study showed that the SP/NK-1R axis was upregulated in PSC patients.^4^ IHC for NK-1R demonstrated that liver tissues from PSC patients showed higher immunoreactivity for NK-1R than those of control (normal) patients (Figure 5A), indicating that SP/NK-1R is upregulated in the liver of PSC patients as expected. SP is synthesized from its precursor TAC1 and deactivated by MME. We have previously demonstrated that TAC1 expression is upregulated in the liver samples of PSC patients.^4^ To determine NK-1R expression in specific hepatic cells, we performed double-staining IF in 2D primary cells. NK-1R and CK-19 (cholangiocyte marker) were co-localized, and immunoreactivity of NK-1R was elevated in CK-19^+^ cholangiocytes in PSC patients (Figure 5B), indicating that cholangiocytes express NK-1R and its expression levels are upregulated during PSC. NK-1R expression in desmin^+^ HSCs was also observed (Figure 5C). These findings indicate that the SP/NK-1R axis is associated with PSC.

**Figure 5:**
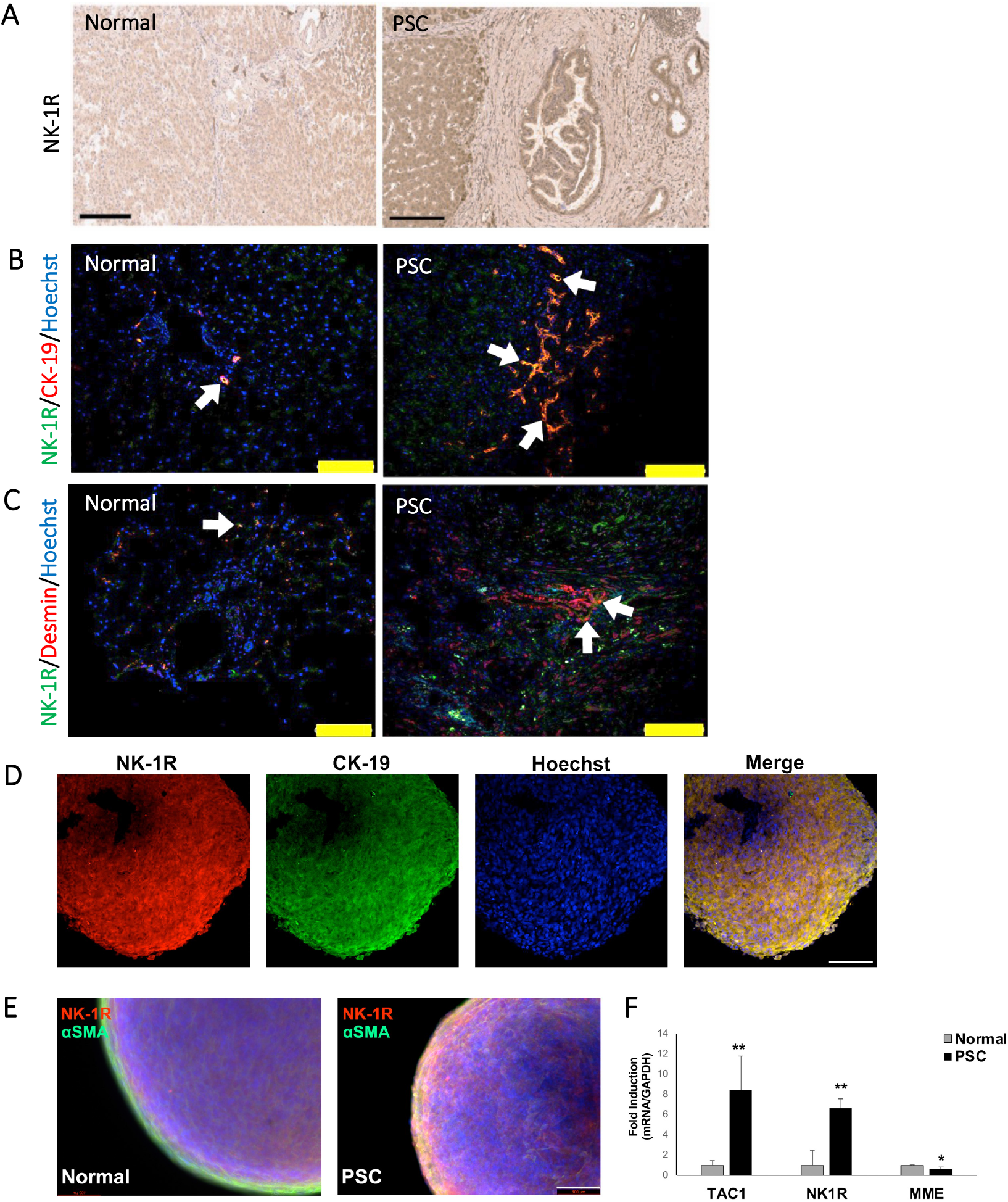
Neuroendocrine phenotype of PSC cell lines and 3D-CHOs. **A**. IHC for human liver sections to detect NK-1R. Scale bar: 200 µm. Representative images of 20x. **B**. NK-1R expression in primary human cholangiocytes. The cell nucleus was stained with Hoechst (in blue). IF to detect NK-1R (green) in human CK-19^+^ cholangiocytes (red) of normal and PSC patients. Arrows point to the area of interest. Scale bar: 100 µm. Representative images 40x. **C**. NK-1R expression in primary human HSCs from normal and PSC patients. Arrows point to the area of interest. Scale bar: 100 µm. Representative images 40x. **D**. Detection of protein expression by IF in live cholangiocyte organoids. NK-1R (red) and CK-19 (green) expression was detected by double staining IF. **E**. Representative IF co-staining image for NK-1R (red) and αSMA (in green) in normal and PSC organoids. **F**. Expression levels of TAC1, NK-1R, and MME on normal and PSC cholangiocyte organoids (n=3 per experiment).

**Figure 6:**
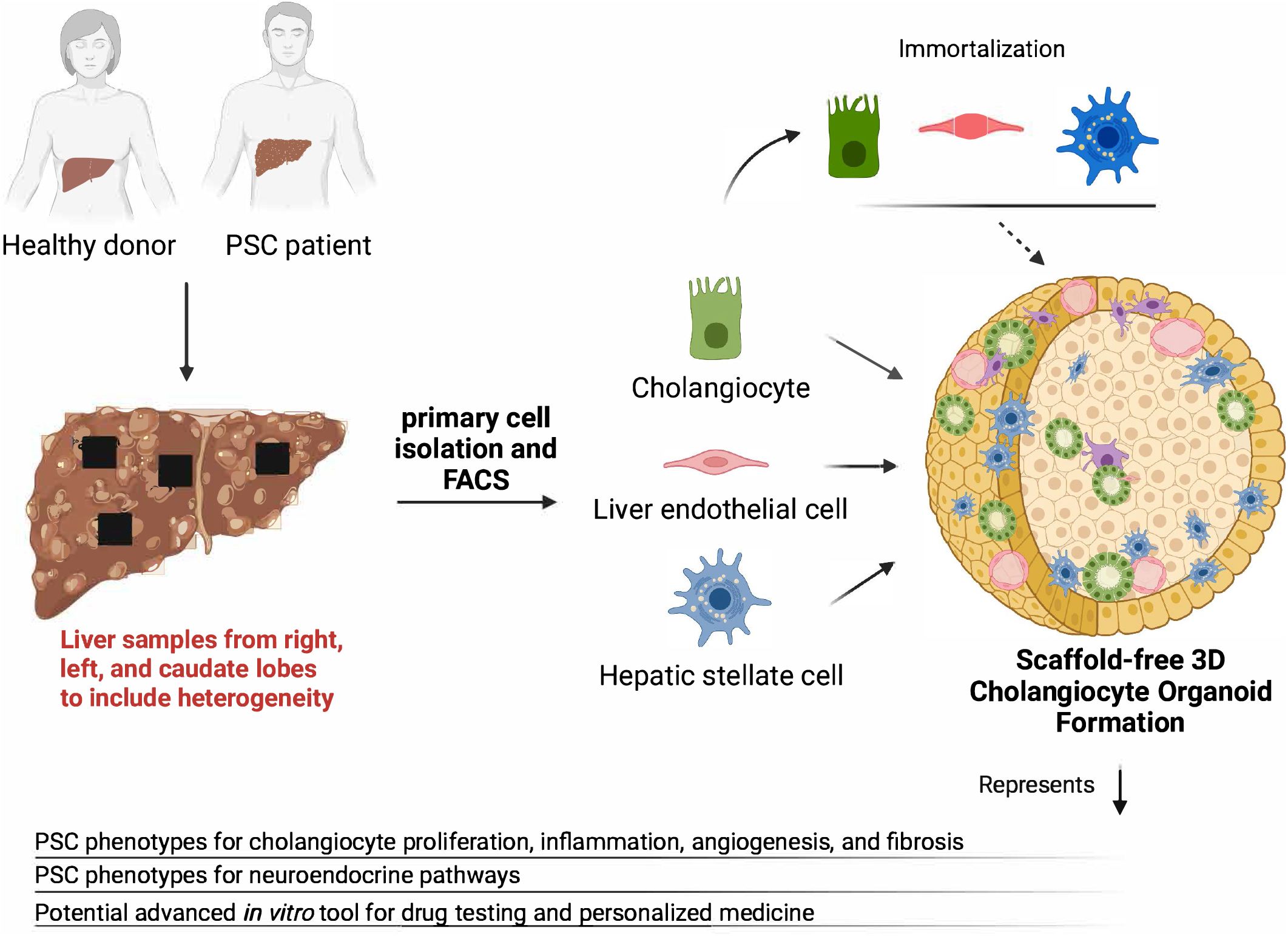
Schematic illustration of the formation of scaffold-free 3D cholangiocyte organoids which mimic the PSC 3D microenvironment. This schematic illustration shows that primary human liver cells can be isolated from normal and PSC patients from multiple liver samples, such as right lobe, left lobe, and caudate lobe to include heterogeneity of the disease. These cells can be immortalized if needed. After organoid formation including multiple liver cell lines without any gel matrix (scaffold), these scaffold-free 3D cholangiocyte organoids can mimic normal and PSC liver microenvironment and phenotypes. These viable 3D cholangiocyte organoids can be used for (i) specific pathway assessment, such as substance P/neurokinin-1 receptor pathway or (ii) drug testing and personalized medicine in PSC since genotype responses will be able to be tested (e.g., Aprepitant treatment, an FDA-approved neurokinin-1 receptor antagonist) (illustration created using biorender.com)

Next, we performed IF in 3D-CHOs to examine NK-1R protein expression in PSC organoids. Double staining for CK-19 and NK-1R showed the strong co-localization of NK-1R in CK-19^+^ cholangiocytes (Figure 5D). Moreover, double staining for NK-1R and αSMA showed the co-localization of NK-1R in αSMA^+^ HSCs is minimal in normal 3D-CHOs; however, the expression of NK-1R in αSMA^+^ HSCs was significantly upregulated in PSC 3D-CHOs (Figure 5E), which phenocopied the NK-1R expression upregulation in HSCs of PSC liver tissue (Figure 5C). Further, the qRT-PCR analysis demonstrated a profound upregulation of TAC1 and NK-1R and downregulation of MME in 3-cell PSC organoids (Figure 5F), indicating SP/NK-1R signaling axis is elevated in PSC organoids similar to the liver tissues from PSC patients, confirming again the neuroendocrine phenotype of 3D-CHOs similar to PSC patients.

## DISCUSSION

It has been previously shown that the isolation of high-purity cholangiocytes from PSC patients was feasible, and those PSC cholangiocytes demonstrated increased senescence-associated signaling compared to normal cholangiocytes in 2D cultures.^9^ Loarca et al.^11^ also showed that 3D organoids could be formed from PSC cholangiocytes, which they named ‘cholangioids’. More recently, bile-derived organoids developed from patients with PSC^10^, and liver-tissue-derived cholangiocyte organoids from PSC patients^19^ have been reported with encouraging outcomes to recapitulate inflammatory immune profile^10^, and cholangiopathy-associated cell death.^19^ However, recent studies^10,19^ and other studies using PSC organoid models^8,11,12,20^ used single cell-cholangiocyte-based organoids. Although 3D organoids have provided many novel findings to better understand and mimic human PSC *in vitro* compared to 2D cell cultures, the main limitation of all the above-mentioned studies was the lack of liver microenvironment in which cell-to-cell crosstalk is the hallmark of disease processes.

To our knowledge, no previous study has shown (i) the isolation and full characterization of HSC and LEC cell lines from human PSC livers, or (ii) the development of 3-cell (cholangiocytes+HSCs+LECs) 3D-CHOs from ‘primary’ human cells viable for up to 1 month. In the current study, we demonstrated, for the first time, the development of viable scaffold-free human PSC organoids from multiple liver cells, rather than stem cell-derived hepatocyte-like or cholangiocyte-like cells or single-cell, cholangiocyte-derived organoids. The inclusion of multiple liver cell lines, such as HSCs, and LECs together with cholangiocytes, created a liver niche with the inclusion of cell-to-cell crosstalk.^21^

Our primary cell-based organoid system has several advantages, including primary liver cells, such as HSCs and LECs isolated from biopsy-proven PSC livers at the time of explant during liver transplantation. This allows the evaluation of PSC phenotypes, not only in cholangiocytes but also in other supporting liver cells. By using primary cells directly from PSC livers, we are able to measure genotypes and personalized responses to certain therapeutic pathways (e.g., SP/NK-1R axis, Figure 5), and will be able to test potential drug candidates. In fact, interpersonal genotype differences cannot be studied in a small animal model or in commercially available stem cell-derived cholangiocyte-like or other liver cell-like cells/organoids since the drug response will always be against the ‘single’ donor of those stem cells. The use of multiple primary liver cells, such as LECs and HSCs together with cholangiocytes creates a true 3D microenvironment with cell-to-cell crosstalk and therefore represents a niche for the studied disease. Most importantly, our organoid system does not use any supporting gel matrix (e.g., Matrigel) allowing it to be scaffold-free and biomaterial-free. Therefore, it does not include any confounders in genomic and/or proteomic analysis, as it happens with the use of Matrigel derived from mouse sarcoma cells.^6,7^

Most previous organoid studies, including PSC organoids, used Matrigel as the extracellular matrix (ECM) component.^8,10-12^,^19, 20^ In fact, ring-like ductal structures were commonly observed in the bipotent ductal organoid culturing condition with Matrigel as a supporting matrix.^18,20^ In the current study, the formation of ring-like ductal structures in 3-cell 3D-CHOs indicates that HSCs and LECs provide signaling cues to orchestrate human cholangiocyte organoid formation and ECM support under the scaffold-free 3D culturing conditions (Figure 3D). Moreover, we established cholangiocyte organoids with fully differentiated cell types that mimic cellular interaction in the portal area, including cholangiocytes, HSCs, and LECs. Interestingly, our scaffold-free method enables us to evaluate organoid ECM secretion leading to liver fibrosis *in vitro*.

Consistently, hepatic cells orchestrate responses against liver damage via the intercellular crosstalk.^21^ The functions of cholangiocytes and HSCs are closely related during cholestatic liver injury, and inhibition of cholangiocyte proliferation decreased ductular reaction, HSC activation, and fibrogenesis.^22,23^ *In vivo*, damaged cholangiocytes secrete elevated levels of profibrogenic cytokine, such as TGFβ1, which activates HSCs leading to ECM secretion and liver fibrosis.^22,23^ Interestingly, we observed an increased expression of fibrosis marker genes, such as Acta2, COL1A1, and desmin and angiogenesis marker genes, such as PECAM, CDH5, and vWF in PSC organoids compared to normal organoids, which indicates that our 3D-CHOs mimic the abnormal activation of HSC and angiogenesis that are often observed in patients during PSC progression. In fact, several assays confirmed PSC phenotypes in our scaffold-free 3D-CHOs with activated cholangiocytes, increased fibrosis, increased angiogenesis, and elevated inflammation (Figures 3, 4, and 5).

Previously, we have demonstrated *in vivo* that serum secretin levels were elevated in PSC patients and BDL and *Mdr2*^-/-^ (PSC model)^22,24^, and expression levels of SR in cholangiocytes were upregulated during cholestatic liver injury in BDL rats.^25^ In the present study, our 3-cell PSC organoids showed similar outcomes to *in vivo* models with increased secretin and SR levels compared to normal cholangiocyte organoids confirming again similar gene expression profiles and phenotypic validity. In addition, we further studied and confirmed the neuroendocrine phenotypic changes in 3D-CHOs with the demonstration of co-localization of upregulated NK-1R with CK-19^+^ cholangiocytes and with αSMA^+^ HSCs, as well as upregulated NK-1R and TAC1 expression, and downregulated MME expression.

There were some limitations in our study. First, we did not check the fibrosis level of each explanted PSC liver sample listed in Table 1 even though our clinical observation of cirrhosis was different in each sample, as shown in Figures 1A, 1B, and 1C. Hence, we compared grouped PSC 3D-CHOs with grouped normal 3D-CHOs. Second, due to the main objective of the study which was the development of scaffold-free multi-cellular 3D-CHOs, we did not compare 3D-CHOs one by one to each other including patient characteristics and differences as listed in Table 1. We believe that these comparisons can be done in future studies when the functionality or drug responses will be tested and compared in different PSC patients.

In conclusion, we have successfully developed scaffold-free multi-cellular 3D cholangiocyte organoids from primary cells to study the pathophysiological mechanisms driving PSC progression. Although there are possibilities that a prolonged culturing period and cell immortalization could cause the alternation of cellular properties, we demonstrated that our 3D-CHO system phenocopied many of the hallmark features of PSC disease, such as biliary inflammation, fibrosis, and angiogenesis. By further refining the culturing conditions, including primary cholangiocytes, HSCs, LECs, and other relevant liver cell types, such as hepatocytes and Kupffer cells, our cholangiocyte/liver organoid system can be used as an advanced novel *in vitro* model to study the PSC disease and test potential therapeutic strategies for the treatment of PSC.^26^

## Abbreviations

αSMA: smooth muscle α-actin
BDL: bile duct ligation
CFTR: cystic fibrosis transmembrane conductance regulator
COL1A1: Collagen Type 1 alpha 1
DMEM: Dulbecco’s Modified Eagle Medium
DMSO: dimethyl sulfoxide
ECM: extracellular matrix
EGF: epidermal growth factor
EMEM: Eagle’s Minimum Essential Medium
EpCAM: epithelial cellular adhesion molecule
FACS: flow cytometry
FBS: fetal bovine serum
GFAP: glial fibrillary acidic protein
HSC: hepatic stellate cell
IF: immunofluorescence
IHC: immunohistochemistry
LEC: liver endothelial cell
Mdr2^-/-^: multidrug resistant-2-knockout
MME: membrane metalloendopeptidase
NK-1R: neurokinin-1 receptor
NPC: non-parenchymal cell
PBS: phosphate-buffered saline
P-gp: P-glycoprotein
POSTN: periostin
PSC: primary sclerosing cholangitis
qRT-PCR: Quantitative PCR
Rh123: Rhodamine-123
SP: substance P
SR: secretin receptor
TAC1: tachykinin precursor 1
VE-Cadherin: vascular endothelial cadherin
vWF: von Willebrand factor
3D: three dimensional
3D-CHO: 3D cholangiocyte organoid

## Authorship

Conceptualization, methodology, software, formal analysis, investigation, data curation, visualization, writing – original draft (WZ, BE); Methodology, investigation, writing –review & editing (AI, YP, PL, ACN, KL, LK, KS); Resources, supervision, writing – review & editing (GA, BE); Conceptualization, resources, funding acquisition, project administration, writing – review & editing (SG, HF, GA, BE); Conceptualization, resources, supervision, funding acquisition, project administration, writing – review & editing (GA, BE). All authors critically reviewed and participated in the writing of the paper. All authors approved the final version.

## REFERENCES

1. Lazaridis KN, LaRusso NF. Primary Sclerosing Cholangitis. N Engl J Med 2016; 375(12): 1161–1170.

2. Ikenaga N, Liu SB, Sverdlov DY, Yoshida S, Nasser I, Ke Q, Kang PM, Popov Y. A new Mdr2(-/-) mouse model of sclerosing cholangitis with rapid fibrosis progression, early-onset portal hypertension, and liver cancer. Am J Pathol 2015; 185(2): 325–334.

3. Glaser S, Gaudio E, Renzi A, Mancinelli R, Ueno Y, Venter J, White M, Kopriva S, Chiasson V, DeMorrow S, Francis H, Meng F, Marzioni M, Franchitto A, Alvaro D, Supowit S, DiPette DJ, Onori P, Alpini G. Knockout of the neurokinin-1 receptor reduces cholangiocyte proliferation in bile duct-ligated mice. Am J Physiol Gastrointest Liver Physiol 2011; 301(2): G297–305.

4. Wan Y, Meng F, Wu N, Zhou T, Venter J, Francis H, Kennedy L, Glaser T, Bernuzzi F, Invernizzi P, Glaser S, Huang Q, Alpini G. Substance P increases liver fibrosis by differential changes in senescence of cholangiocytes and hepatic stellate cells. Hepatology 2017; 66(2): 528–541.

5. Li M, Izpisua Belmonte JC. Organoids - Preclinical Models of Human Disease. N Engl J Med 2019; 380(6): 569–579.

6. Kleinman HK, Martin GR. Matrigel: basement membrane matrix with biological activity. Semin Cancer Biol 2005; 15(5): 378–386.

7. Hughes CS, Postovit LM, Lajoie GA. Matrigel: a complex protein mixture required for optimal growth of cell culture. Proteomics 2010; 10(9): 1886–90.

8. Shiota J, Samuelson LC, Razumilava N. Hepatobiliary Organoids and Their Applications for Studies of Liver Health and Disease: Are We There Yet? Hepatology 2021; 74(4): 2251–2263.

9. Tabibian JH, Trussoni CE, O’Hara SP, Splinter PL, Heimback JK, LaRusso NF. Characterization of cultured cholangiocytes isolated from livers of patients with primary sclerosing cholangitis. Lab Invest 2014; 94(10): 1126–33.

10. Soroka CJ, Assis DN, Alrabadi LS, Roberts S, Cusack L, Jaffe AB, Boyer JL. Bile-derived organoids from patients with primary sclerosing cholangitis recapitulate their inflammatory immune profile. Hepatology 2019; 70(3): 871–882

11. Loarca L, De Assuncao TM, Jalan-Sakrikar N, Bronk S, Krishnan A, Hunag B, Morton L, Trussoni C, Marcano Bonilla L, Kruger E, O’Hara S, Splinter P, Shi G, Pisarello MJL, Gores GJ, Huebert RC, LaRusso NF. Development and characterization of cholangioids from normal and diseased human cholangiocytes as an in vitro model to study primary sclerosing cholangitis. Lab Invest 2017; 97(11): 1385–1396.

12. Nuciforo S, Heim MH. Organoids to model liver disease. JHEP Rep 2021; 3(1): 100198.

13. Mederacke I, Dapito DH, Affo S, Uchinami H, Schwabe RF. High-yield and high-purity isolation of hepatic stellate cells from normal and fibrotic mouse livers. Nat Protoc 2015; 10(2): 305–315.

14. Zhang Y, Zeng Z, Zhao J, Li Dali, Liu M, Wang X. Measurement of Rhodamine 123 in Three-Dimensional Organoids: A Novel Model for P-Glycoprotein Inhibitor Screening. Basic Clin Pharmacol Toxicol 2016; 119(4): 349–352.

15. Wang Z, Faria J, van der Laan LJW, Penning LC, Masereeuw R, Spee B. Human cholangiocytes form a polarized and functional bile duct on hollow fiber membranes. Front Bioeng Biotechnol 2022; 10: 868857.

16. Kadioglu, O., et al., Interactions of human P-glycoprotein transport substrates and inhibitors at the drug binding domain: Functional and molecular docking analyses. Biochem Pharmacol, 2016. 104: p. 42–51.

17. Singh, R., et al., In vitro effects of standardized extract of Bacopa monniera and its five individual active constituents on human P-glycoprotein activity. Xenobiotica, 2015. 45(8): p. 741–9.

18. Sato K, Zhang W, Safarikia S, Isidan A, Chen AM, Li P, Francis H, Kennedy L, Baiocchi L, Alvaro D, Glaser S, Ekser B, Alpini G. Organoids and Spheroids as Models for Studying Cholestatic Liver Injury and Cholangiocarcinoma. Hepatology 2021; 74(1): 491–502.

19. Shi S, Verstegen MMA, Roest H, Ardisasmita AI, Cao W, Roos FJM, de Ruiter PE, Niemeijer M, Pan Q, Ijzermans JNM, van der Laan Ljw. Recapitulating cholangiopathy-associated necroptotic cell death in vitro using human cholangiocyte organoids. Cell Mol Gastroenterol Hepatol 2022; 13(2): 541–564

20. Wu F, Wu D, Ren Y, Huang Y, Feng B, Zhao N, Zhang T, Chen X, Chen S, Xu A. Generation of hepatobiliary organoids from human induced pluripotent stem cells. J Hepatol 2019; 70(6): 1145–1158.

21. Sato K, Kennedy L, Liangpunsakul S, Kusumanchi P, Yang Z, Meng F, Glaser S, Francis H, Alpini G. Intercellular communication between hepatic cells in liver diseases. Int J Mol Sci 2019; 20(9): 2180.

22. Wu N, Meng F, Invernizzi P, Bernuzzi F, Venter J, Standeford H, Onori P, Marzioni M, Alvaro D, Franchitto A, Gaudio E, Glaser S, Alpini G. The secretin/secretin receptor axis modulates liver fibrosis through changes in transforming growth factor-b1 biliary secretion in mice. Hepatology 2016; 64(3): 865–879.

23. Wu N, Meng F, Zhou T, Venter J, Giang TK, Kyritsi K, Wu C, Alvaro D, Onori P, Mancinelli R, Gaudio E, Francis H, Alpini G, Glaser S, Franchitto A. The secretin/secretin receptor axis modulates ductular reaction and liver fibrosis through changes in transforming growth factor-beta1-mediated biliary senescence. Am J Pathol 2018; 188(10): 2264–2280.

24. Wu, N, Baiocchi L, Zhou T, Kennedy L, Ceci L, Meng F, Sato K, Wu C, Ekser B, Kyritsi K, Kundu D, Chen L, Meadows V, Franchitto A, Alvaro D, Onori P, Gaudio E, Lenci I, Francis H, Glaser S, Alpini G. Functional role of the secretin/secretin receptor signaling during cholestatic liver injury. Hepatology 2020; 72(6): 2219–2227.

25. Alpini G, Ulrich CD, Philips JO, Pham LD, Miller LJ, LaRusso NF. Upregulation of secretin receptor gene expression in rat cholangiocytes after bile duct ligation. Am J Physiol 1994; 266: G922–928.

26. Sato K, Glaser S, Kennedy L, Liangpunsakul S, Meng F, Francis H, Alpini G. Preclinical insights into cholangiopathies: disease modeling and emerging therapeutic targets. Expert Opin Ther Targets 2019; 23(6): 461–472.

